# EEG Signatures of Auditory Distraction: Neural Responses to Spectral Novelty in Real-World Soundscapes

**DOI:** 10.1101/2025.04.14.648656

**Authors:** Silvia Korte, Thorge Haupt, Martin G. Bleichner

**Affiliations:** Neurophysiology of Everyday Life Group, Department of Psychology, University of Oldenburg, Oldenburg, Germany; Research Center for Neurosensory Science, University of Oldenburg, Oldenburg, Germany

## Abstract

In everyday life, ambient sounds can disrupt our concentration, interfere with task performance, and contribute to mental fatigue. Even when not actively attended to, salient or changing sounds in the environment can involuntarily divert attention. Understanding how the brain responds to these real-world auditory distractions is essential for evaluating the cognitive consequences of environmental noise. In this study, we recorded electroencephalography (EEG) while participants performed different tasks during prolonged exposure to a complex urban soundscape. We identified naturally occurring, acoustically salient events and analyzed the corresponding event-related potentials (ERPs). Auditory spectral novelty reliably elicited a P3a response (250–350 ms), reflecting robust attentional capture by novel environmental sounds. In contrast, the Reorienting Negativity (RON) window (450–600 ms) showed no consistent modulation, possibly due to the continuous and largely behaviorally irrelevant nature of the soundscape. Performance in a behavioral task was briefly disrupted following novel sounds, underscoring the functional impact of attentional capture. Noise sensitivity, measured via the Weinstein Noise Sensitivity Scale (WNSS; 1978), was not associated with ERP amplitudes. Together, these findings demonstrate that the P3a component provides a stable neural marker of attentional shifts in naturalistic contexts and highlight the utility of spectral novelty detection as a tool for investigating auditory attention outside the laboratory.

## 1 Introduction

### 1.1 The Importance of Irrelevant Sounds

In our increasingly noisy environments (Asdrubali, 2014), managing auditory attention is crucial for cognitive performance, well-being, and brain health. While in complex acoustic environments we need to focus on relevant sounds, such as a conversation or an approaching vehicle, we also need to suppress or ignore irrelevant sounds (Ahveninen et al., 2011; Choi, Rajaram, Varghese, & Shinn-Cunningham, 2013; Fritz, Elhilali, & Shamma, 2005; Hillyard, Hink, Schwent, & Picton, 1973; Schwartz & David, 2018; Shinn-Cunningham & Best, 2008). This process is cognitively demanding and can cause mental fatigue over time (Saremi et al., 2008). Understanding how the brain copes with persistent background noise and how individual differences shape this process (Kjellberg, Landström, Tesarz, Söderberg, & Akerlund, 1996), is essential for designing quieter workspaces, improving public spaces, and mitigating the negative effects of noise pollution.

Studies on auditory distractibility propose a three-phase model of distraction processing (Escera, Alho, Schröger, & Winkler, 2000; Getzmann, Arnau, Gajewski, & Wascher, 2024; Wetzel & Schröger, 2014). First, a change in the acoustic environment is detected by an automatized deviance detection mechanism. This change may be yielded by a violation of expectations and leads to a pre-attentive processing of the detected deviant. The second phase is an involuntary shift in attention to the deviating event, leading to a deduction of attentional resources from task-relevant information. Lastly, a voluntary re-orienting away from the deviant sound is done and expectations about the soundscape are updated.

Suppressing or ignoring irrelevant sounds can involve both bottom-up and top-down mechanisms. While some distracting sounds may never reach conscious awareness due to automatic filtering at early sensory stages (e.g., Boutros and Belger (1999)), others require effortful suppression that drains cognitive resources (Bidet-Caulet et al., 2007; Schwartz & David, 2018). As this ongoing effort accumulates, it can contribute to increased mental fatigue and even feelings of annoyance that are associated with psychological, psychiatric, and cardiovascular diseases (Basner et al., 2014). However, auditory distraction is not experienced uniformly across individuals; some are more susceptible to attentional capture by novel sounds, while others exhibit stronger cognitive control mechanisms to suppress distraction (Kjellberg et al., 1996; Shepherd, Hautus, Lee, & Mulgrew, 2016). Investigating how these individual differences shape neural responses to irrelevant sounds could provide insight into why some listeners are more affected by noise than others.

Given these challenges, it is important to consider how well-established methods in auditory neuroscience translate to everyday environments. In particular, the tools we use to study attention must account for the complexity and unpredictability of real-world soundscapes.

### 1.2 From Controlled Stimuli to Real-World Soundscapes

Traditional event-related potential (ERP) paradigms have typically relied on short, repetitive, and highly controlled auditory stimuli. While these designs have been instrumental in advancing our understanding of auditory processing (e.g., Hillyard et al. (1973); Spong, Haider, and Lindsley (1965)), they offer limited insight into how attention operates in complex, everyday environments. Real-world listening is defined by dynamic, overlapping streams of sound that unfold continuously over time. This raises questions about the extent to which findings from laboratory paradigms generalize to natural listening situations.

In response, some studies have moved toward more ecologically valid designs by embedding naturalistic sounds, such as speech or environmental noises, into continuous audio streams (e.g., Rosenkranz, Cetin, Uslar, and Bleichner (2023); Straetmans, Holtze, Debener, Jaeger, and Mirkovic (2021)). These “middle-ground” approaches preserve some aspects of real-world acoustics while retaining the experimental structure required for meaningful EEG interpretation. Our previous work followed this rationale, demonstrating that neural responses to repeated naturalistic stimuli are shaped by factors like task complexity and personal relevance (Korte, Jaeger, Rosenkranz, & Bleichner, 2024). However, such studies still rely on artificial structuring of the soundscape.

Fully natural soundscapes, such as a busy street recording, present a further challenge. They are dense, unpredictable, and do not contain any experimenter-controlled stimuli. While listeners are adept at noticing salient changes (Hicks & McDermott, 2024), such as a sudden honk or tram bell, identifying these moments objectively is non-trivial. Real-world acoustic events often overlap or are masked by background noise, much like how subtle instruments in polyphonic music can be masked by louder ones (Müller, 2021). Their salience depends on spectral, temporal, and contextual factors (Lavie, 2005; Müller, 2021) and are not necessarily represented in the waveform of the audio.

To overcome these limitations, the current study utilizes a spectral novelty detection algorithm (Müller, 2021) to identify perceptually salient auditory events directly from the natural soundscape. It eliminates the need for artificially embedded stimuli, allowing us to examine neural responses to naturally occurring, context-defined sounds in complex acoustic environments.

ERPs serve as a key tool for this investigation, capturing the brain’s time-locked responses to discrete events. Within the framework of the three-phase model of auditory distraction (Escera et al., 2000; Getzmann et al., 2024; Wetzel & Schröger, 2014), ERPs provide markers for each stage of the process: the initial detection of deviance is reflected by the Mismatch Negativity (MMN), a negative deflection occurring around 100 ms at central and prefrontal sites. In the second phase the involuntary shift of attention is marked by the P3a component, a positive deflection peaking around 300 ms at the frontal midline. Lastly, the voluntary re-orienting away from the distractor is indexed by the Reorienting Negativity (RON), a frontocentral negativity occurring between 400 and 600 ms after stimulus onset. By examining these components in response to naturally occurring auditory changes, we aim to assess how the brain manages distraction in a real-world context.

## 2 Methods

### 2.1 Participants

The present study builds on the dataset reported in Korte et al. (2024). In total, 30 individuals underwent audiometric screening (pure-tone audiometry). 23 participants (13 female, 10 male) met the eligibility criterion of having hearing thresholds of at least 20 dB HL at octave frequencies from 250 Hz to 8 kHz and were included in the final sample. Participants were between 21 and 37 years old (mean: 25.57, SD: 3.48), right-handed, had normal or corrected-to-normal vision, and reported no history of neurological, psychiatric, or psychological conditions. All participants provided written informed consent and received monetary compensation for their participation.

### 2.2 Procedure

Prior to EEG data acquisition, participants completed the Weinstein Noise Sensitivity Scale (WNSS; Weinstein 1978), a 21-item inventory designed to assess individual differences in noise sensitivity. The questionnaire asks participants to rate their agreement with statements related to noise (e.g., “I wouldn’t mind living on a noisy street if the apartment I had was nice.”) on a 6-point Likert scale ranging from “strongly disagree” to “strongly agree.” The total WNSS score reflects a participant’s general sensitivity to noise, with higher scores indicating greater susceptibility to noise-related annoyance and distraction.

Afterwards, participants completed six blocks of EEG recordings, each lasting 15 to 45 minutes, totaling approximately 3.5 hours of recording data. Participants could take self-determined breaks between blocks. The experimental design alternated between passive listening blocks, where participants were instructed to disregard the soundscape, and active listening conditions, where they responded to specific auditory events.

### 2.3 Paradigm

All parts of the paradigm, apart from the transcription task, were presented using the Psychophysics Toolbox extension (Brainard 1997; Kleiner, Brainard, and Pelli 2007; Pelli 1997, Version: 3) on MATLAB 2021b.

#### 2.3.1 Auditory Stimuli

For our analysis in this paper, we only included blocks in which the pre-recorded soundscape of a busy city street^1^ was played. It consisted of a variety of ambient sounds, typical of an urban area (e.g., streetcars, motorcycles or incomprehensible speech). The street scenario had a total length of 2 hours and 21 minutes, from which we took four segments of 45 minutes each. These segments had a short overlap, since the original sound file was not long enough to cover three non-overlapping hours. The sequence of segments was randomized across participants.

In the original study, additional auditory stimuli (church bells) were added to the street soundscape. However, for the analysis presented in this paper, only the urban soundscape without additional auditory cues was considered.

#### 2.3.2 Non-auditory Task

Participants engaged in one of two non-auditory tasks, depending on the experimental block. The first was a visual search task using detailed hidden object pictures, similar to the well-known “Where’s Waldo?/Where’s Wally?” game. Participants searched for specific objects within complex illustrated scenes and selected them using the mouse. The number of targets was deliberately set high to ensure continuous engagement throughout the block.

The second task was a transcription task that resembled simple office work. Taken from the citizen science project “World Architecture Unlocked” on “Zooniverse”, this task required participants to transcribe handwritten details from architectural photographs^2^. Participants categorized information such as city names, architects, or building names. No prior architectural knowledge was necessary, but the task was sufficiently complex to require sustained attention. Participants were encouraged to use search engines and online maps to verify and categorize the transcribed information. The task involved reading, typing, using the mouse, and researching, making it a suitable approximation of realistic office-based cognitive tasks, while providing a higher task complexity than the hidden-object picture task. In the original experiment, the street soundscape was played during four separate blocks, each combined with either the visual search task or the transcription task. These blocks were not designed for the current analyses but were selected post hoc because they provided extended, ecologically valid exposure to a complex auditory environment while participants were engaged in non-auditory activities.

#### 2.3.3 Experimental Blocks

The experiment consisted of three phases, divided into 4 consecutive blocks, as illustrated in Figure 1. The passive phase A comprised 2 blocks, while the active phase and the passive phase B each included one block.

**Figure 1:**
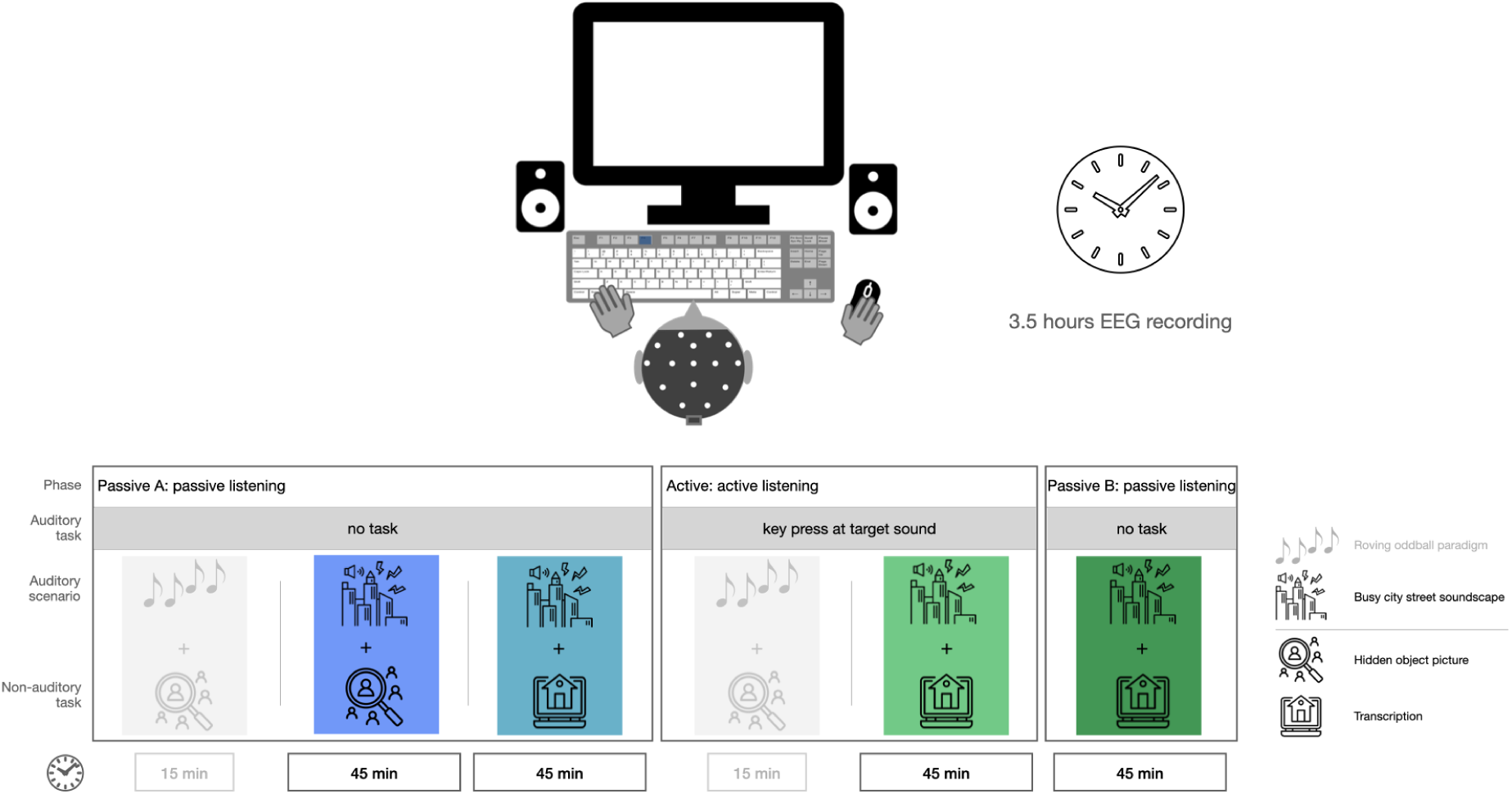
Overview of experimental blocks. Order of the blocks is chosen to ensure naivety concerning target sounds in the passive phase A. Gray shaded blocks are not considered in this work and are only displayed for the sake of completeness.

During the experiment, participants were either instructed that the soundscape was irrelevant or that they had to detect a specific target sound (church bell). This manipulation was intended to draw participants’ attention to the overall soundscape. In the passive listening conditions, participants were told that background sounds would be present but were irrelevant to their task and could be ignored. In the active listening condition, participants were required to detect and respond to the target sound by pressing the F4 key. The response key was chosen to avoid interference with the non-auditory task, where the keyboard was also used. All other sounds in the soundscape were not behaviorally relevant and did not require a response.

The block structure and auditory manipulations were part of a broader dataset that has been described in detail in our previous work (Korte et al., 2024). The present study focuses on a subset of the experimental blocks from that study. The structure for the present study was as follows:

- Block 1 (Passive listening A, Passive Phase A): A 45-minute sequence of the street soundscape was played while participants performed the hidden-object picture task (HOP), without responding to the sounds.
- Block 2 (Passive listening A, Passive Phase A): Identical to Block 1, but participants performed the transcription task.
- Block 3 (Active listening, Active Phase): Identical to Block 2, except participants now responded to the target sounds (church bell) in the street soundscape.
- Block 4 (Passive listening B, Passive Phase B): Identical to Block 2. However, participants were instructed to ignore the previously relevant church bell chimes again.

The order of blocks was fixed for all participants to ensure that they remained naive to the target sound during the passive listening phase A and to preserve the mental representation of the target sound in the passive listening B phase. Randomizing the order would have disrupted the intended transition between conditions, particularly in the final block. This design allowed for a consistent progression across participants.

### 2.4 Data Acquisition

#### 2.4.1 Description of Lab Setup

Participants were seated in a soundproof recording booth at a desk equipped with a screen (Samsung, SyncMaster P2470). A keyboard and a mouse were placed on the desk for task input and target response. Auditory stimuli were played via two free-field loudspeakers (Sirocco S30, Cambridge Audio, London, United Kingdom) positioned at ear level, at a 45-degree angle to the left and right with a distance of approximately 0.5 m. The soundscape was played at an average volume of 51 dB(A) using loudspeakers. Event markers for auditory stimuli and task events were generated using the Lab Streaming Layer (LSL) library^3^. Keyboard input was logged using LSL-compatible key capture software^4^. The Lab Recorder software^5^ ensured synchronized data recording of the EEG data, the event markers and the keyboard capture in .xdf format. Files were organized using the Brain Imaging Data Structure (BIDS) format (Gorgolewski et al., 2016) with the EEG data extension (Pernet et al., 2019).

#### 2.4.2 EEG system

EEG data were collected using a 24-channel EEG cap (EasyCap GmbH, Hersching Germany) with passive Ag/AgCl electrodes (channel positions: Fp1, Fp2, F7, Fz, F8, FC1, FC2, C3, Cz, C4, T7, T8, CP5, CP5, CP1, CPz, CP2, CP6, TP9, TP10, P3, Pz, P4, O1, O2). The mobile cap setup, with fewer electrodes than typical lab systems, was chosen for participant comfort during the extended recording sessions and was well tolerated, even during breaks. A mobile amplifier (SMARTING MOBI, mBrainTrain, Belgrade, Serbia) was attached to the EEG cap, allowing participants a more natural sitting position compared to a wired EEG system. Gyroscope data from the amplifier were recorded to track head movements. Data were transmitted via Bluetooth to a desktop computer using a BlueSoleil dongle. EEG and gyroscope data were streamed to LSL via the SMARTING Streamer software (v3.4.3; mBrainTrain, Belgrade, Serbia) and recorded at a sampling rate of 250 Hz using Lab Recorder.

#### 2.4.3 Measurement Procedure

Before data collection, electrode sites were cleaned with 70 % alcohol and abrasive gel (Abralyt HiCl, Easycap GmbH, Germany). Electrode gel was applied to maintain impedances below 10 kΩ and impedances were monitored throughout the session. If signal quality dropped, individual electrodes were re-gelled between blocks. Re-gelling was rare, typically affecting one or two electrodes, and no full cap removal was required. Given the experiment’s length, participants were allowed an extended lunch break, scheduled to avoid interference with experimental manipulation (active phase and passive phase B).

### 2.5 Data Analysis

All analyses were conducted in MATLAB 2021b using the EEGLAB toolbox (Delorme and Makeig 2004; version: 2021.1).

#### 2.5.1 Behavioral Data

To investigate whether sound events interfered with participants’ typing behavior, we analyzed the time interval between consecutive keystrokes. Specifically, we examined whether inter-keystroke intervals (IKIs) were longer when a sound onset occurred between two keystrokes, compared to intervals without an intervening sound. The IKI was defined as the time between the first and the second keystroke. If the sound was perceived as distracting, the second keystroke was expected to be delayed, resulting in a longer interval. To ensure comparability, control intervals were selected immediately prior to sound events. Only IKIs between 200 and 600 ms were retained to exclude implausibly short latencies and those unlikely to reflect continuous typing. For each participant and condition, mean IKI values were computed separately for sound and no-sound intervals.

#### 2.5.2 Audio Processing and Feature Extraction

We were interested in how listeners perceive complex street soundscapes, particularly how auditory events influence attention and neural responses. In an initial test run, two human listeners manually annotated perceptually salient events in the soundscape. Their annotations confirmed that distinct auditory objects were identifiable and corresponded to measurable brain responses when used to timelock EEG data. However, manual annotation is both time-consuming and highly variable, depending on factors such as headphone use, attentional state, and individual listener differences.

To address this, we applied spectral novelty analysis (Müller, 2021), an algorithmic method for detecting salient sound events based on abrupt changes in spectral content—especially in higher frequencies, where transient sounds are more easily distinguished from background noise. This approach enables reproducible event detection in naturalistic soundscapes without relying on manual annotations or artificial stimuli.

We implemented the method using the open-source MATLAB functions `spectral novelty.m` and `simp peak.m` (available at github.com/ThorgeHaupt/Audionovelty). Audio recordings of the street scenes (sample rate: 44.1 kHz) were converted to mono by averaging stereo channels. Each signal was transformed into the time-frequency domain using a Short-Time Fourier Transform (STFT; Hanning window size: 882 samples, hop size: 441 samples). The resulting magnitude spectrogram was logarithmically compressed (*γ* = 10) to enhance perceptually relevant spectral variations (Figure 2, second plot from top, left panel).

**Figure 2:**
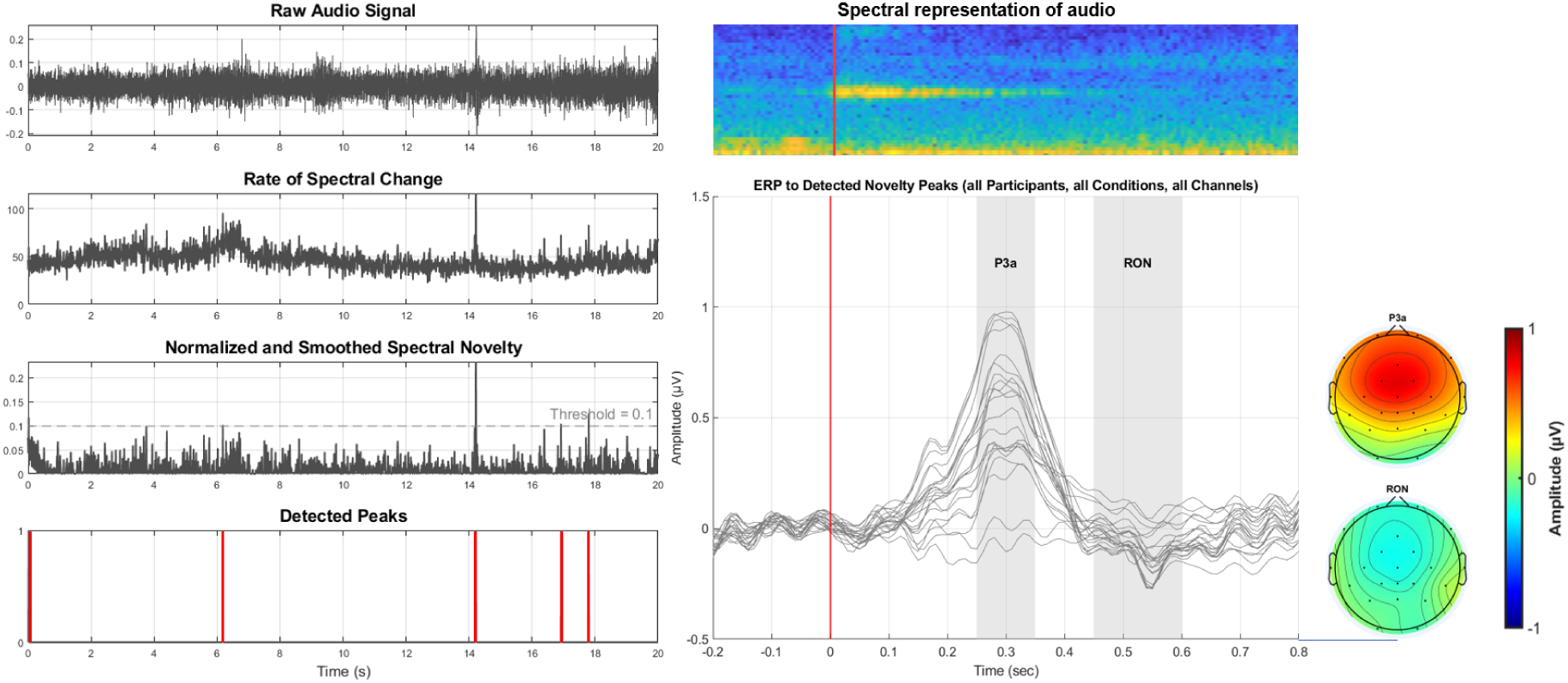
Left: Example for spectral novelty decomposition on a snippet of 20 seconds. The raw audio (top plot) is first transformed into the time-frequency domain, from which a spectral change is computed (second plot). Afterwards, this spectral change is normalized and smoothed (third plot). Lastly, spectral peaks are identified based on a fixed threshold, resulting in a binary vector of peaks (bottom plot); Top right: spectral representation of an example novelty event. Bottom right: Resulting ERP and topographies of P3a- and RON-time-window. Each trace represents one EEG channel.

To highlight spectral changes, a first-order derivative across time frames was computed. Negative values were set to zero, and a local average over a 0.5 s window was subtracted to reduce noise and emphasize meaningful fluctuations (Figure 2, third plot from top, left panel). The resulting novelty function was normalized and resampled to 100 Hz to match the temporal resolution required for further EEG analysis.

Sound onsets were defined as local maxima in the novelty function that exceeded neighboring values and a fixed threshold of 0.1. This resulted in a binary peak vector marking moments of salient acoustic change (Figure 2, bottom plot, left panel). These peak markers were used to time-lock EEG analyses to naturally occurring auditory events in the urban soundscape.

#### 2.5.3 EEG Data

The EEG was preprocessed as described in Korte et al. (2024). To ensure clarity, we briefly summarize the processing steps here. The data were first filtered between 1 and 40 Hz (default settings of the `pop eegfiltnew` function). Next, bad channels were identified, using the clean artifacts function of EEGLAB with the default settings for `channel crit maxbad time` and subsequently stored for later interpolation. The data were then segmented into 1-second windows. Artifact rejection was performed on these windows based on a probability threshold of ±3 SD from the mean, which helped optimize independent component analysis (ICA) training. All data were then combined to compute ICA weights using the `runica` function in EEGLAB with the extended training mode. The ICA weights were applied to the raw EEG data.

Artifact rejection was performed using the ICLabel algorithm (Pion-Tonachini, Kreutz-Delgado, & Makeig, 2019), where components classified with ≥ 80 % probability as artifacts (e.g., eye blinks, muscle activity, heartbeats) were removed. Additionally, a manual inspection was conducted to account for possible misclassifications, as the ICLabel algorithm is primarily optimized for stationary datasets with minimal movement, whereas our setup allowed participants a degree of mobility. On average, 8 out of 24 components were removed per participant (min = 4, max = 11).

Following ICA-based artifact removal, the EEG data were further processed by applying a low-pass filter at 20 Hz and a high-pass filter at 0.5 Hz. Any previously identified bad channels were interpolated (mean = 1.4, min = 0, max = 5), and the data were re-referenced to the average of electrodes Tp9 and Tp10, which never had to be interpolated. EEG data were resampled to 100 Hz to match the temporal resolution of the spectral novelty function.

Events corresponding to the onset of spectral novelty peaks were identified, and their latencies were mapped to the EEG time series. Epochs were extracted from -0.2 to 0.8 seconds relative to sound onset. If an epoch extended beyond the available data range, zero-padding was applied to maintain uniform epoch length across trials. Baseline correction was performed using a pre-stimulus interval from 0.2 to 0 seconds, subtracting the mean baseline activity from each epoch.

To assess differences in neural processing of the soundscapes under different listening conditions, we computed a grand average ERP and topographic maps for each block. Additionally, we investigated the relationship between neural responses and spectral novelty. To assess whether the magnitude of the neural response depends on the degree of novelty of a given sound, consistent with expectations based on Downar, Crawley, Mikulis, and Davis (2002), where several brain areas showed sensitivity to stimulus novelty, epochs were sorted according to their novelty score and assigned to 20 equally sized bins, sorted in ascending order of spectral novelty. Bins were equalized (where necessary) by excluding excess trials randomly (with a fixed random seed for reproducibility). Artifactual epochs were removed after binning, with a probability criterion of ±3 SD from the mean. A 50 % overlap between bins was applied to ensure smoother transitions between novelty levels.

To investigate ERP responses, we focused on four frontocentral electrodes (Fz, FC1, FC2, Cz), selected based on previous literature emphasizing their sensitivity to components within the three-phase model of auditory distraction (Escera et al., 2000; Getzmann et al., 2024; Wetzel & Schröger, 2014). For each participant and condition, ERPs were averaged across these electrodes and across trials. To enable cross-subject comparison, we further computed grand-average ERPs per novelty bin across participants.

The P3a component was analyzed as the mean amplitude in the 250–350 ms time window postonset, while the RON component was assessed in the 450–600 ms window. These windows were selected based on visual inspection of peak deflections in the grand-average waveforms and align with typical latencies reported in the literature (Escera et al., 2000; Getzmann et al., 2024; Wetzel & Schröger, 2014). We did not include the MMN in our analysis, as its elicitation typically depends on a structured sequence of frequent standard and infrequent deviant stimuli (Näätänen, Paavilainen, Titinen, Jiang, & Alho, 1993), which introduce a violation of regular auditory patterns. Since our continuous, real-world street soundscape is highly dynamic by nature, no such pattern of consistent auditory regularities exist. Thus, our soundscape is not suited to investigate the MMN component.

#### 2.5.4 Statistical Analysis

##### Weinstein Noise Sensitivity Scale (WNSS)

In an exploratory analysis, we examined whether individual differences in noise sensitivity predict neural responses to auditory novelty. We conducted Pearson’s correlation analyses between WNSS scores and individual ERP amplitudes averaged over all conditions and as a mean of the selected frontocentral channels. Specifically, we tested the relationship between WNSS scores and mean ERP amplitudes in two key time windows: the P3a window (250–350 ms) and the RON window (450–600 ms). The normality of the WNSS scores and ERP amplitudes was confirmed using the Shapiro-Wilk test (WNSS: p = 0.451, P3a: p = 0.366, RON: p = 0.160), justifying the use of Pearson’s correlation. Correlations were computed separately for the P3a and RON amplitudes, with significance levels set at *p <* 0.05.

##### Behavioral data

Statistical analysis was performed using the Wilcoxon signed-rank test for paired samples to compare typing speed between uninterrupted and interrupted typing within each experimental condition. This non-parametric test was chosen due to deviations from normality in the data distribution, as confirmed by the Shapiro-Wilk test. The Wilcoxon signed-rank test was applied separately for each condition to determine whether the presence of auditory interruptions significantly affected typing speed.

To account for multiple comparisons, p-values were adjusted using the Benjamini-Hochberg False Discovery Rate (FDR) correction. This method controls the expected proportion of false positives while maintaining statistical power.

To assess whether the magnitude of behavioral disruption (i.e., the difference in IKIs between interrupted and uninterrupted typing) differed between conditions, we conducted a Friedman test. This non-parametric equivalent of a repeated-measures ANOVA is suitable for comparing more than two related samples when the data may not follow a normal distribution. The Friedman test was applied to per-subject difference scores across all three experimental conditions that contained the transcription task.

Statistical analyses were conducted in MATLAB R2021b using the `signrank.m` function for Wilcoxon tests and the `mafdr.m` function for FDR correction.

##### EEG data

To assess differences in ERP amplitudes across conditions, we conducted Wilcoxon signed-rank tests, a non-parametric paired test, comparing mean ERP amplitudes within the time windows of interest for the P3a and RON.

We performed four pairwise comparisons, motivated by the study’s design:

- Passive A + HOP vs. Passive A + Transcription, to test whether the type of non-auditory task modulates ERP amplitudes under passive listening conditions.
- Passive A + Transcription vs. Active + Transcription, to examine whether directing attention to the soundscape in the active condition influences ERP amplitudes.
- Active + Transcription vs. Passive B + Transcription, to determine whether ERP amplitudes remain modulated after the active phase or return to passive A levels.
- Passive A + HOP vs. Passive B + Transcription, to evaluate whether ERP amplitudes in the Passive B Phase differ from those in the Passive A Phase.

To correct for multiple comparisons, we applied False Discovery Rate (FDR) correction using the Benjamini-Hochberg procedure.

To examine the influence of novelty intensity at the single-trial level, we used linear mixed-effects models (LMMs) with novelty score as a continuous predictor. Separate LMMs were fitted for the P3a and RON time windows. The models included fixed effects for novelty score and condition and a random intercept for subject to account for within-subject variability (see Supplement Figure 9). We compared two models per time window using a Likelihood Ratio Test (LRT):

1. Full model: EEG Amplitude ∼ Novelty Score + Condition + (1 | Subject)
2. Simpler model: EEG Amplitude ∼ Novelty Score + (1 | Subject)

Model selection was based on Akaike Information Criterion (AIC) and the likelihood ratio test (LRT) to determine whether including condition improved model fit. The models were implemented using the `fitlme.m` function in Matlab.

## 3 Results

### 3.1 Overall ERP Responses to Spectral Novelty Peaks

In a first step, we investigated the overall ERPs per condition, time-locked to all identified spectral novelty peaks. Figure 3 displays the time-series from -0.2 to 0.8 seconds relative to peak onsets, with corresponding topographical representations for two distinct time windows: 0.25 to 0.35 seconds, corresponding the the P3a and 0.45 to 0.60 seconds, corresponding to the RON.

**Figure 3:**
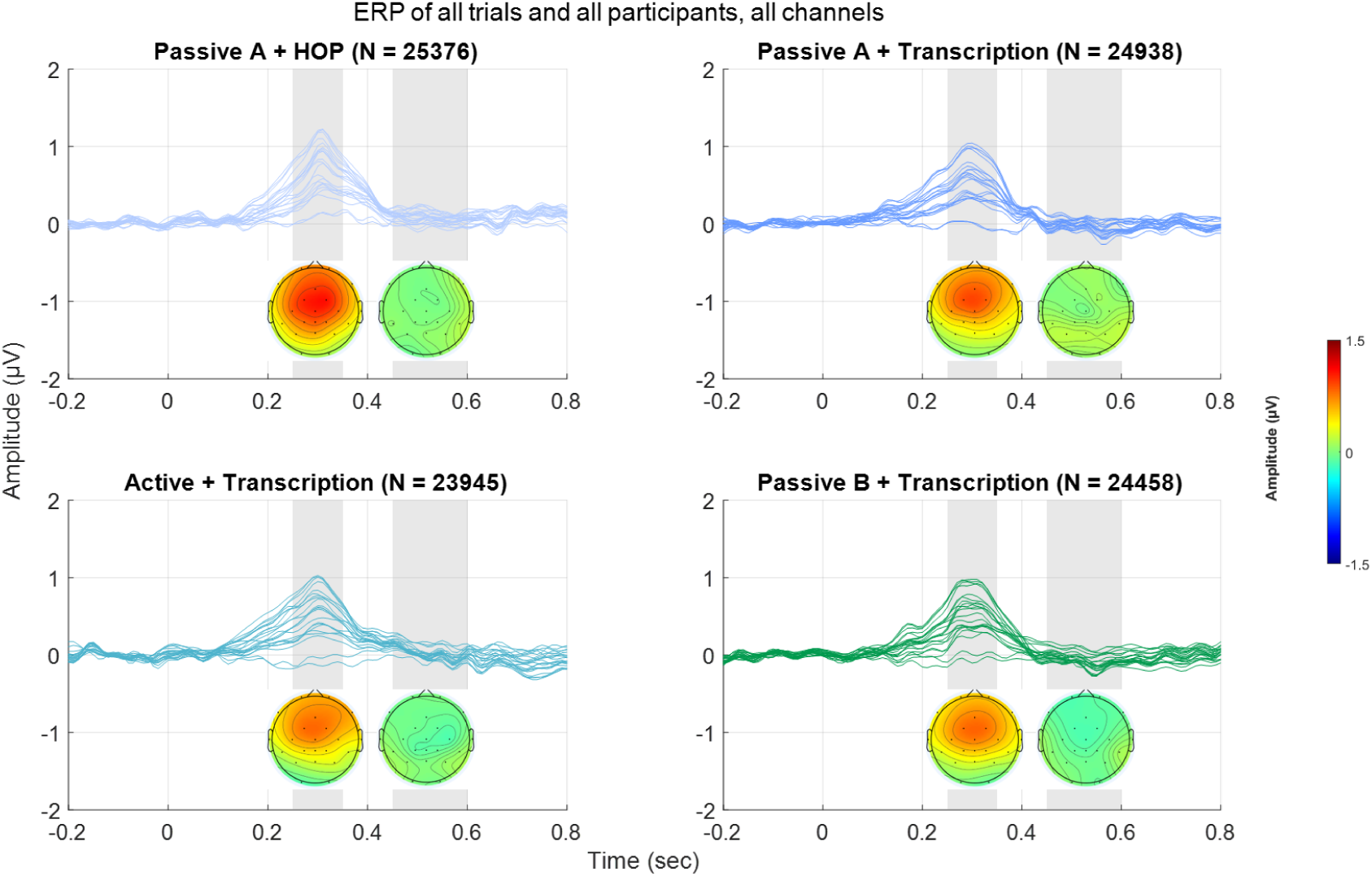
ERPs time-locked to spectral novelty peaks across all participants and trials, visualized as butterfly plots (i.e., each trace represents one EEG channel). Gray shaded area in the time-series plots represent the time-windows for the topographies (0.25 - 0.35 sec and 0.45 - 0.60 sec). N refers to the number of trials included in the average. Top left: Passive listening A condition while participants engaged in the hidden-object picture task (HOP). Top right: passive listening A condition while participants engaged in the transcription task. Bottom left: Active listening condition while participants engaged in the transcription task. Bottom right: Passive listening B while participants engaged in the transcription task.

A distinct positive deflection at approximately 300 ms post-onset can be observed across all conditions, with the strongest amplitude around frontocentral electrodes. This pattern is consistent with the expected characteristics of the P3a component. The response is most pronounced in the passive listening A condition with the hidden-object picture task (HOP), followed by the passive listening A condition with the transcription task. The amplitude of this peak appears slightly reduced in the active listening condition and lowest in the passive listening B condition.

In contrast, we did not observe a pronounced negative deflection in the expected RON time window (450–600 ms). While there are slight amplitude variations across conditions, the expected negativity is not clearly present. This suggests that the reorienting process might be weaker or less reliably elicited in the given experimental context. The summary statistics for these analyses are presented in Table 1.

**Table 1:** Summary statistics for the grand ERPs per condition in time windows of interest and for frontocentral channels (mean of Fz, FC1, FC2, Cz)

| Condition | P3a: mean ± SD (μV) | RON: mean ± SD (μV) |
| --- | --- | --- |
| Passive A + HOP | 0.963 ± 0.514 | 0.000 ± 0.445 |
| Passive A + Transcription | 0.867 ± 0.541 | -0.077 ± 0.506 |
| Active + Transcription | 0.859 ± 0.537 | -0.034 ± 0.425 |
| Passive B + Transcription | 0.898 ± 0.612 | -0.096 ± 0.479 |

### 3.2 Condition Effects on ERP Amplitudes

#### P3a Time Window (250 - 350 ms)

A Wilcoxon signed-rank test was conducted to compare ERP amplitudes across conditions. None of the pairwise comparisons showed a significant difference in ERP amplitude before or after correction for multiple comparisons (all p-values *>* .05). Specifically, comparisons between Passive A + HOP and Passive A + Transcription (W = 165, p = 0.2113, FDR-corrected p = 0.8453), Passive A + Transcription and Active + Transcription (W = 127, p = 0.9870), Active + Transcription and Passive B + Transcription (W = 126, p = 0.9870), and Passive A + HOP and Passive B + Transcription (W = 133, p = 0.8329, FDR-corrected p = 0.9870) all yielded non-significant results.

These findings indicate that neither task engagement nor listening condition (passive vs. active) led to significant changes in the P3a time window.

#### RON Time Window (450 - 600 ms)

To investigate voluntary reorientation processes, we examined ERP amplitudes within the 450–600 ms time window. The Wilcoxon signed-rank tests revealed no significant differences between conditions, even before correction for multiple comparisons (all uncorrected p-values > .05). Specifically, comparisons between Passive A + HOP and Passive A + Transcription (W = 147, p = .5057, FDR-corrected p = .6743), Passive A + Transcription and Active + Transcription (W = 103, p = .4455, FDR-corrected p = .6743), Active + Transcription and Passive B + Transcription (W = 138, p = .7089), and Passive A + HOP and Passive B + Transcription (W = 155, p = .3548, FDR-corrected p = .6743) all yielded non-significant results.

These findings suggest that no robust effects were observed in the RON time window, regardless of task or listening condition.

### 3.3 EEG Responses as a Function of Spectral Novelty

#### P3a component

We observed a systematic increase in EEG amplitude with higher spectral novelty scores (Figure 4), particularly in the Passive A + HOP condition (Figure 6). This trend is visually apparent in the grand average ERPs across novelty bins (Figure 5), where larger P3a amplitudes are observed in bins with higher novelty values.

**Figure 4:**
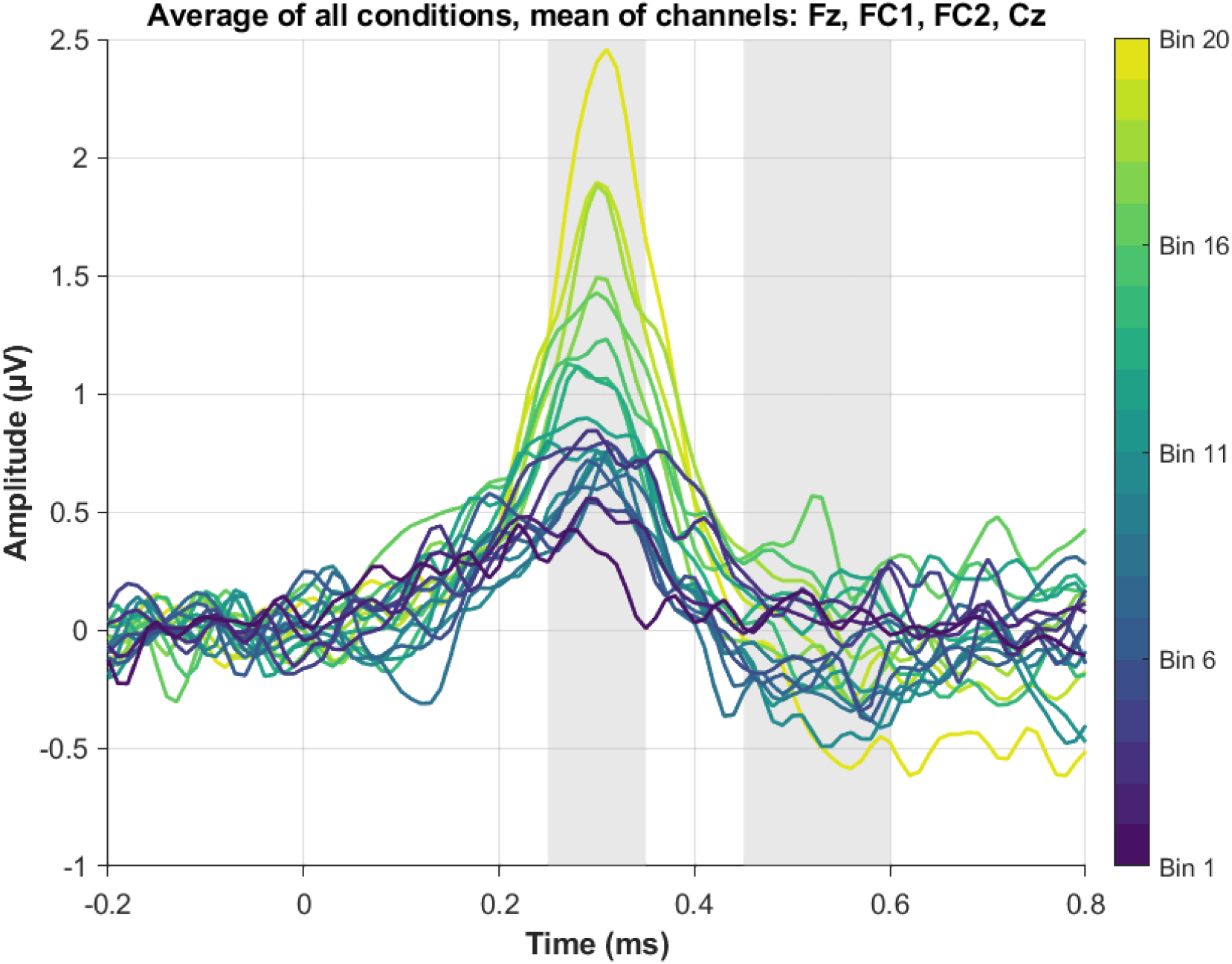
Grand average ERPs across the 20 novelty bins as mean over all conditions. Data were averaged per condition first, then across conditions.

**Figure 5:**
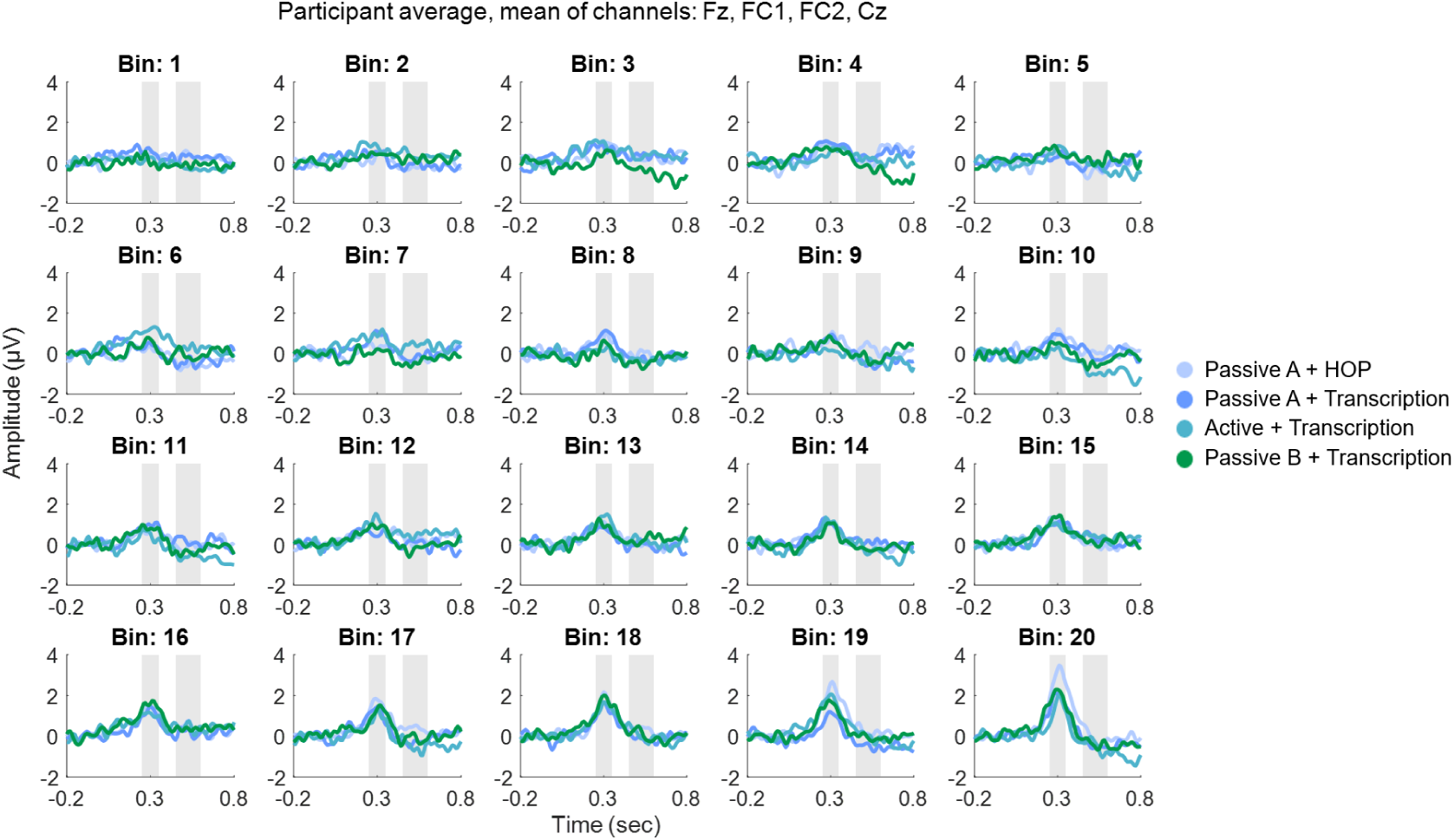
Grand average ERPs across 20 spectral novelty bins for each listening condition. Each subplot represents the ERP response for a specific spectral novelty bin, with Bin 1 corresponding to the lowest novelty values and Bin 20 to the highest. Grey marked windows correspond to the P3a window (early window, 0.25 - 0.35 sec) and the RON window (late window, 0.45 - 0.60 sec).

**Figure 6:**
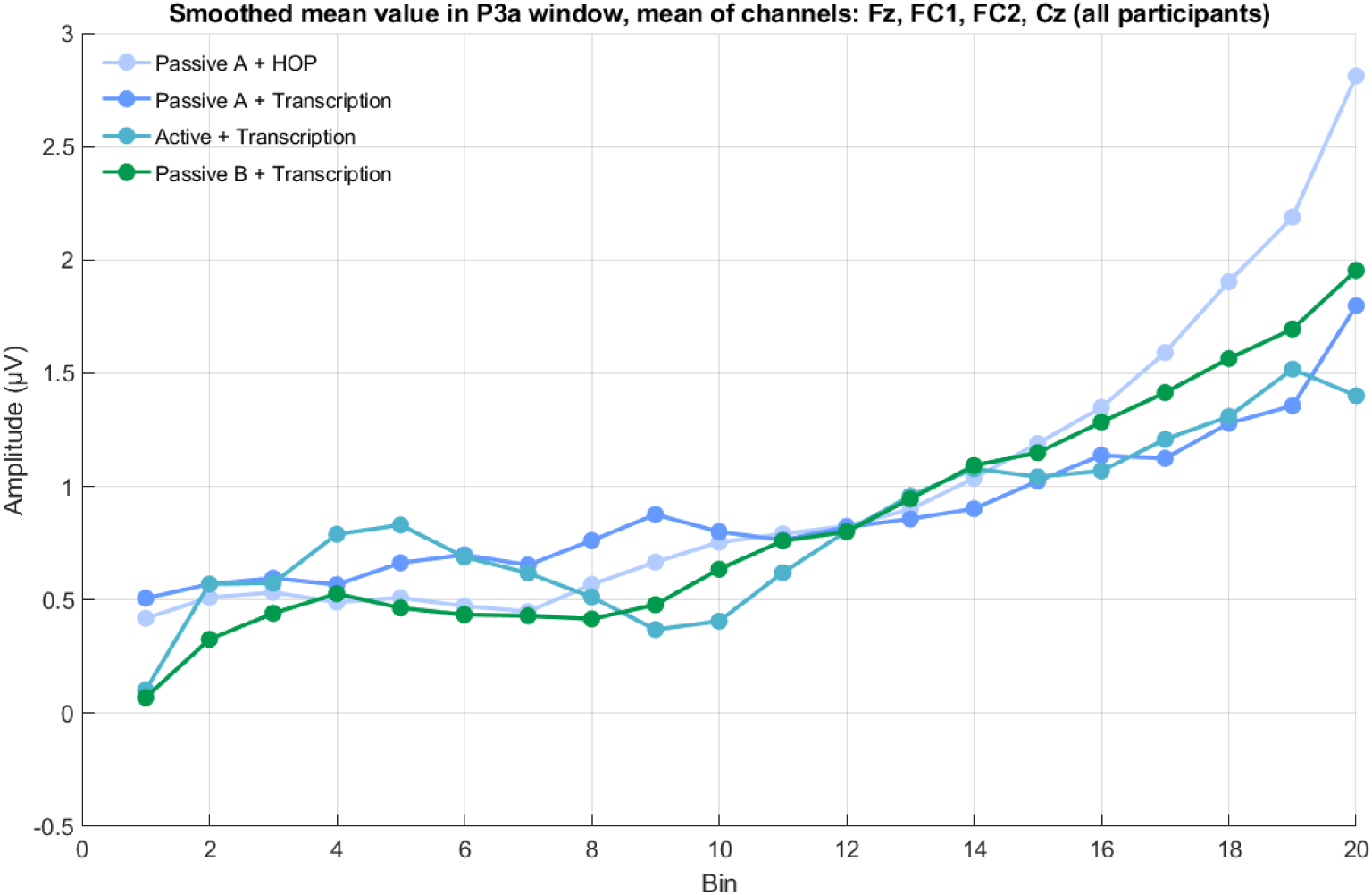
Development of mean ERP amplitudes across frontocentral electrodes (Fz, FC1, FC2, Cz) for each spectral novelty bin.

LMM analysis confirmed a significant effect of novelty score on EEG amplitude (*β* = 4.53, *p < .*001), supporting the observed trend that higher novelty is associated with increased P3a amplitudes. Including “condition” as a predictor did not improve model fit (*χ*^2^(3) = −3.22, *p* = 1), and no reliable pairwise differences between conditions were observed (*p > .*05 for all conditions). While one contrast (Passive A + HOP vs. Active + Transcription) showed a borderline p-value (*p* = .049), this result should be interpreted with caution, as it emerged from a model that did not outperform the simpler one.

These statistical findings align with the visual representation in Figures 4 and 5, where novelty is the primary factor modulating amplitude, and condition differences are subtle. For an illustration of individual participant trajectories across novelty bins, see Supplementary Figure 9, which presents ERP amplitude changes in the P3a time window for each participant, separated by median split. Similarly, the Wilcoxon signed-rank test results confirmed that condition did not significantly influence P3a amplitude.

#### RON component

In contrast to the P3a window, no strong amplitude changes are evident in the RON time window (Figure 5 and 4). The expected negative deflection for the RON component is not clearly present across conditions. LMM analysis confirmed the absence of an effect of novelty (*β* = −0.11, *p* = .823) or condition (*p > .*05 for all pairwise comparisons). Additionally, adding condition to the model did not improve fit (*χ*^2^(3) = 0.00, *p* = 1).

These statistical findings are visually supported by Figure 5 and 4, where no pronounced negativity is evident in the 450–600 ms window. Likewise, Wilcoxon signed-rank tests did not reveal significant differences between conditions, further reinforcing that RON amplitudes are not modulated by spectral novelty or condition.

### 3.4 Behavioral Data

The analysis of IKIs revealed a consistent increase for interrupted typing compared to uninterrupted typing across all three experimental conditions (Figure 7). Descriptive statistics showed that in the Passive A + Transcription condition, the mean IKI was 327.04 ms (SD = 18.17) for uninterrupted typing and increased to 350.03 ms (SD = 24.86) for interrupted typing. In the Active + Transcription condition, the mean IKI was 327.45 ms (SD = 23.48) for uninterrupted typing and 347.99 ms (SD = 37.77) for interrupted typing. Similarly, in the Passive B + Transcription condition, the mean IKI increased from 320.88 ms (SD = 14.30) in the uninterrupted state to 357.03 ms (SD = 49.97) in the interrupted state.

**Figure 7:**
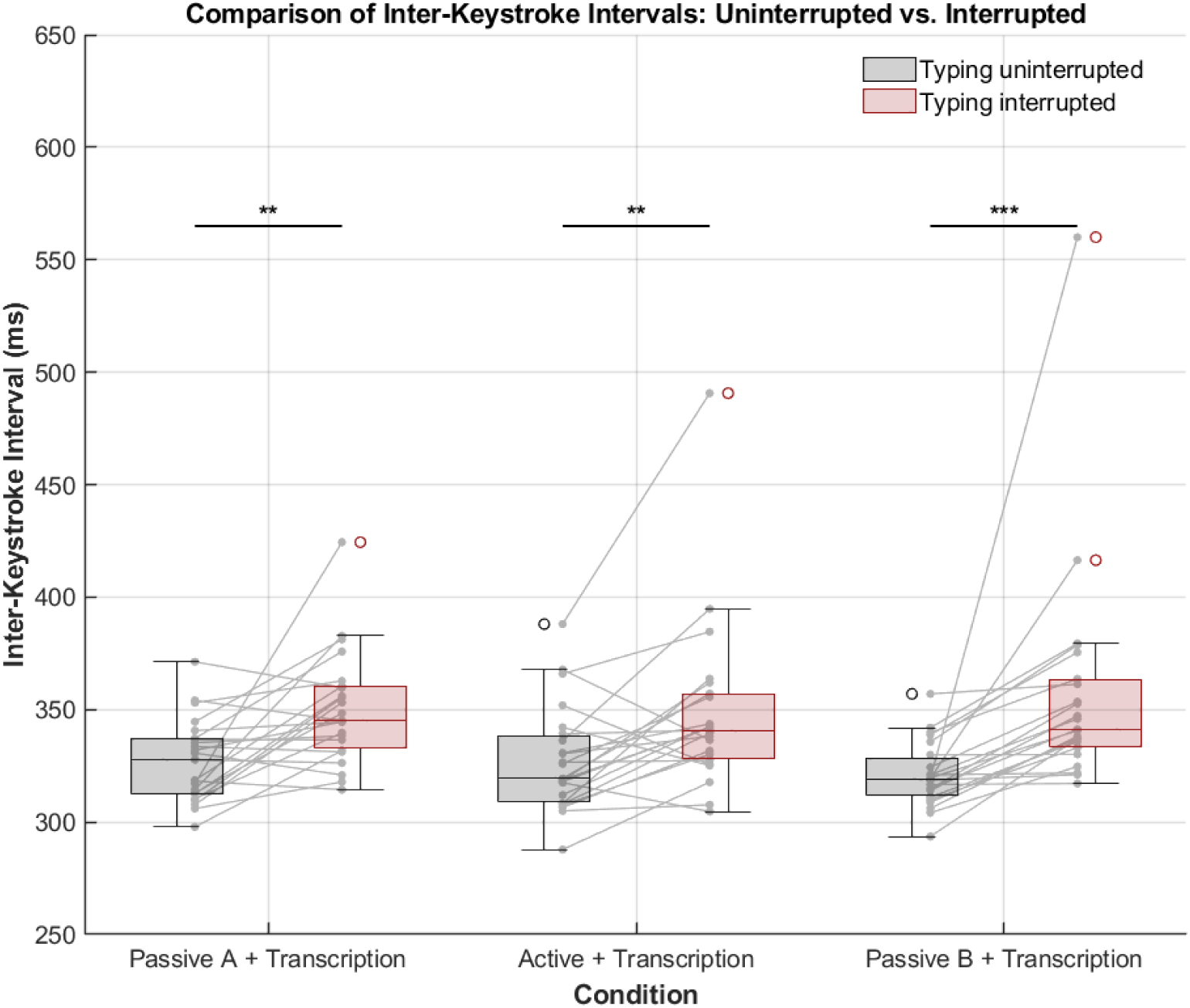
Analysis of inter-keystroke intervals for uninterrupted vs. interrupted typing across conditions. Gray lines connect corresponding values for each participant. Asterisks indicate significance levels (* *p* ≤ 0.05; ** *p* ≤ 0.01; *** *p* ≤ 0.001). Friedman’s test revealed no significant difference between conditions (p = 0.958).

Statistical analysis using the Wilcoxon signed-rank test confirmed that IKIs were significantly larger when typing was interrupted in all conditions. In the Passive A + Transcription condition, the test yielded W = 38, p = 0.0024 (uncorrected), p = 0.0035 (FDR-adjusted). For the Active + Transcription condition, the results were *W* = 42, *p* = 0.0035 (uncorrected), *p* = 0.0035 (FDR-adjusted). The effect was most pronounced in the Passive B + Transcription condition with W = 0, *p <* 0.0001 (uncorrected), *p* = 0.0001 (FDR-adjusted). Furthermore, Friedman test revealed no significant differences in the magnitude of behavioral disruption across conditions, *chi*^2^(2) = 0.09, p = 0.958, indicating that the prolonged IKIs following sound events were comparable across all conditions. These findings indicate that the sound events significantly increased IKIs, suggesting a robust disruptive effect of sound events on typing, irrespective of experimental conditions.

### 3.5 Weinstein Noise Sensitivity Scale (WNSS)

Participants’ noise sensitivity was assessed using the Weinstein Noise Sensitivity Scale (WNSS; Weinstein 1978). The WNSS scores were normally distributed (Shapiro-Wilk test: *p* = 0.451), with a mean score of 3.36 (SD = 0.54, range = 2.43–4.76). These values were previously reported in our earlier study (Korte et al., 2024), which examined the same sample in a different analytical context.

To provide context for these scores, we compared them to the normative value of 3.04 (SD = 0.57), indicating that our sample exhibited slightly higher noise sensitivity on average. However, the observed mean falls within one standard deviation of the norm, suggesting that the noise sensitivity distribution in our sample is broadly comparable to the general population.

To examine whether noise sensitivity was associated with neural responses to auditory novelty, we conducted Pearson’s correlation analyses between WNSS scores and individual ERP amplitudes averaged over all conditions in two key time windows: the P3a window (250–350 ms) and the RON window (450–600 ms).

The correlation analysis revealed no significant relationship between WNSS scores and P3a amplitudes (*r* = −0.006, *p* = 0.980), indicating that noise sensitivity did not predict the degree of attentional capture by novel sounds. A weak negative trend was observed between WNSS scores and RON amplitudes (*r* = −0.192, *p* = 0.393), which would suggest that individuals with higher noise sensitivity may have a less pronounced reorienting response following distraction. However, this trend did not reach statistical significance (see Figure 8). For a visualization of individual ERP waveforms across all participants, see Supplementary Figure 10, which depicts ERPs averaged across all conditions for the selected fronto-central channels.

**Figure 8:**
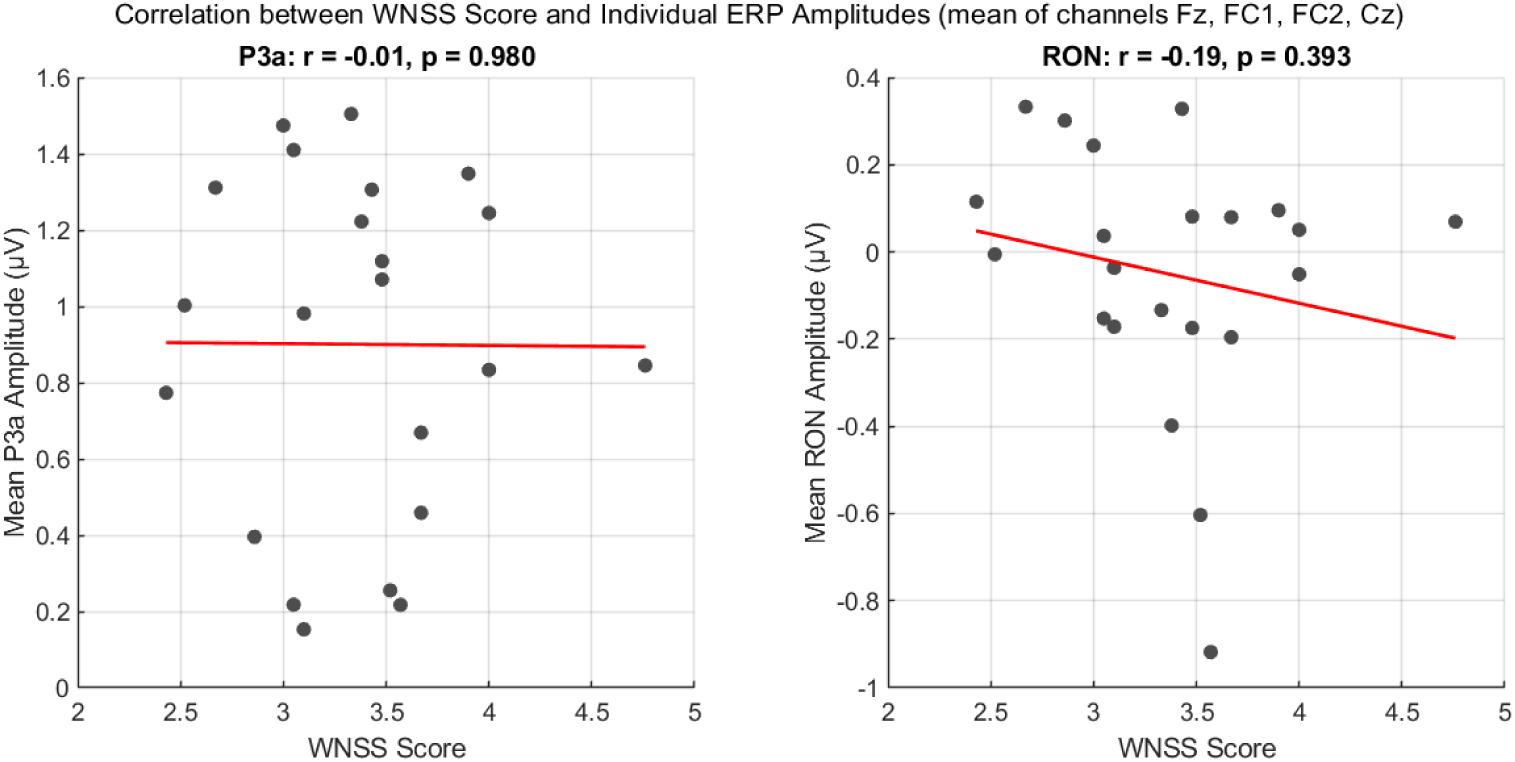
Scatter plots depicting Pearson’s correlation between WNSS scores and mean ERP amplitudes for the P3a (left) and RON (right) components. Each dot represents an individual participant, and red lines indicate the best-fitting linear regression.

## 4 Discussion

This study investigated how the brain processes auditory novelty in a real-world soundscape while participants engaged in different tasks and listening conditions. We found that the intensity of spectral novelty significantly modulated EEG responses, particularly in the P3a time window. This highlights that stronger acoustic changes in the environment evoke more pronounced neural responses. Based on previous research, we expected these neural responses to vary with both novelty intensity and listening engagement. Specifically, we anticipated that active listening would enhance ERP amplitudes compared to passive listening. The contrast between active and passive listening conditions revealed only subtle differences in ERP amplitudes. That is, it did not significantly matter whether participants were passively listening or actively attending to the soundscape. EEG responses to sound events remained robust across all conditions. Below, we discuss the implications of these findings and potential explanations for the observed patterns.

### 4.1 Effect of Spectral Novelty on EEG Responses

The results demonstrated that spectral novelty robustly influenced EEG amplitudes, particularly in the P3a time window. This finding aligns with previous research showing that novel auditory events elicit enhanced neural responses, reflecting attentional capture (Escera et al., 2000; Getzmann et al., 2024; Wetzel & Schröger, 2014).

The increased P3a amplitude we observed with higher novelty values supports the idea that spectral novelty acts as a key trigger for involuntary attentional shifts, which aligns with the second phase of the distraction model. Interestingly, we found no strong modulation in the RON time window, suggesting that participants either did not consistently reorient attention away from novel sounds or that this process was not as pronounced in a naturalistic setting. One possible explanation is that the ongoing soundscape did not provide discrete auditory events that necessitated reorienting, unlike traditional lab paradigms with isolated deviant stimuli. In our setting, novel sounds may not have required context updating or behavioral adaptation, and thus made it unlikely to observe a reorienting response. Additionally, since the background audio was mostly behaviorally irrelevant (even though the active listening condition encouraged attention to the soundscape, where a reorienting response might have been expected), participants may not have engaged in an active re-orienting process. This highlights the need for future research to explore whether reorientation mechanisms are suppressed when auditory distractions occur within continuous, real-world soundscapes rather than discrete, laboratory-controlled paradigms.

Our approach of using spectral novelty provides new insights into how attention dynamically fluctuates in response to environmental sound changes. The observed increase in P3a amplitude with higher novelty (see Figures 4 and 6) suggests that the brain remains highly sensitive to salient changes in the acoustic environment, even when attention is directed elsewhere. This responsiveness likely reflects bottom-up attentional capture by acoustically novel events, consistent with the notion that attention can be involuntarily drawn to unexpected changes in the environment. Furthermore, our findings highlight the robustness of the P3a component, which showed similar amplitude and morphology (i.e., shape, latency) across all conditions. This is in line with previous research (Fallgatter et al., 2000; Korte et al., 2024) and renders the P3a particularly useful for real-world EEG applications, where data might be noisier, and less robust components could be overshadowed by noise. A supplemental figure shows the consistency of this pattern across individual participants (see Supplementary Figure 10). The stability of the P3a across different listening conditions suggests that it may serve as a reliable neural marker for attentional capture in complex auditory environments such as workplaces (Wascher et al., 2023), classrooms (Janssen et al., 2021), or public spaces (Gramann, 2024).

### 4.2 Limited Condition Effects and Potential Explanations

Contrary to our initial hypothesis, listening mode did not significantly alter ERP responses. While the active listening condition was expected to enhance auditory processing compared to passive listening, we found only subtle differences between conditions (see Figure 5 for a visual comparison across bins and conditions). One possible explanation is that attentional resource allocation varied dynamically across tasks, but did not create large enough differences to be reflected in ERPs. The HOP task, being cognitively less demanding than transcription, may have allowed participants to allocate more resources to background sound processing (Sörqvist, Dahlström, Karlsson, & Rönnberg, 2016). This could have facilitated stronger neural responses to auditory novelty, but may not have yielded sufficiently large neural differences to reach statistical significance.

Additionally, habituation effects may have contributed to the condition pattern, particularly since the Passive A + HOP condition was always presented first. This initial exposure to the soundscape may have triggered stronger neural responses, reflecting heightened sensitivity to novel auditory input. While one might expect habituation to continue progressively across all blocks, it is also possible that the largest adjustment occurred early on, during the first encounter with the soundscape. In this view, the strongest novelty-related responses would be limited to the first block, with a relatively stable lower responsiveness in subsequent blocks. Such an early adjustment would be consistent with the deviance detection and attentional shift stages of the three-phase distraction model, which are known to diminish once novelty is no longer perceived as salient.

Another factor to consider is the role of self-generated sounds in the transcription task. Participants’ typing may have introduced competing auditory input that either acoustically masked or perceptually deprioritized the street soundscape. Prior research suggests that self-generated sounds are processed differently from externally generated ones (e.g., Bäß, Jacobsen, and Schröger 2008; Martikainen 2004; Saupe, Widmann, Trujillo-Barreto, and Schröger 2013) and often involve predictive mechanisms that suppress their neural representation. As a result, the soundscape may have been pushed into the perceptual background, both because it was masked by keystrokes and because attentional resources were directed toward the motor task and its auditory consequences. This may have reduced the salience of the background noise, thereby contributing to the weaker ERP responses in conditions involving transcription. Future studies could explore whether minimizing self-generated auditory input alters neural responses to environmental sounds.

### 4.3 Behavioral Impact of Novel Sounds on Task Performance

Beyond the neural effects of spectral novelty, our results also revealed a significant behavioral impact. We observed that IKIs increased in response to novel sounds irrespective of experimental conditions, indicating that distraction effects extend beyond electrophysiological responses (Figure 7). This result aligns with prior research showing that auditory events can momentarily disrupt ongoing cognitive-motor tasks (e.g., Conrad et al. 2012). The slowing of typing speed suggests that involuntary attentional capture, as reflected in the P3a component, translated into measurable performance decrements.

Notably, the behavioral disruption was present across conditions, further supporting the robustness of spectral novelty in capturing attention regardless of task engagement. This aligns with findings from workplace distraction studies, where unpredictable background sounds, such as sudden conversations or environmental noises, reduce productivity in cognitively demanding tasks (Conrad et al., 2012; Kjellberg et al., 1996; Sexton & Helmreich, 2000; Sonnleitner et al., 2014). The observed slowing of typing speed indicates that novel sounds impact task performance in the moments directly following the sound event. Whether this effect extends beyond the immediate keystroke remains to be investigated. Future research should explore whether the distraction persists over time or diminishes with continued exposure. Additionally, while our findings point to a general effect of novelty on behavior, it remains unclear whether varying levels of novelty intensity produce graded effects on task performance. Although this analysis was not feasible in the current dataset due to the limited number of keypress instances per participant, it represents a promising avenue for future investigation.

### 4.4 Individual Differences in Noise Sensitivity and Attentional Modulation

While previous research has highlighted variability in how individuals respond to auditory distraction (Kjellberg et al., 1996; Shepherd et al., 2016), our analysis found no significant correlation between noise sensitivity (WNSS scores) and ERP amplitudes. This suggests that while noise sensitivity may influence subjective experiences of distraction, it does not necessarily translate to differences in early neural responses to background sounds. However, this does not preclude noise sensitivity from affecting later cognitive or behavioral stages of distraction processing.

One possibility is that noise sensitivity exerts its influence beyond early attentional capture, modulating higher-order cognitive and emotional responses to auditory distractions rather than automatic neural responses measured by ERPs. For instance, individuals with higher noise sensitivity may not show stronger P3a responses but may still perceive background noise as more disruptive, leading to greater cognitive fatigue, annoyance, or task disengagement over time.

It is also worth noting that our sample exhibited relatively average noise sensitivity scores, with no extreme outliers. This restricted variability may have limited the ability to detect significant correlations with ERP amplitudes. Future studies should aim to include individuals across a broader range of sensitivity levels to better assess whether more noise-sensitive individuals show distinct neural or behavioral response patterns.

Given this, future research should also explore whether noise sensitivity influences behavioral performance, subjective distraction ratings, or physiological measures (e.g., autonomic responses such as heart rate variability or skin conductance), which may better reflect individual differences in real-world auditory distraction. Additionally, non-linear effects should be considered, as highly noise-sensitive individuals may show disproportionate responses compared to those with lower sensitivity (Kliuchko, Heinonen-Guzejev, Vuust, Tervaniemi, & Brattico, 2016).

### 4.5 Implications for Real-World Auditory Attention Research

Our study highlights the utility of a spectral novelty-detection approach for identifying salient auditory events within a continuous, real-world soundscape. Unlike traditional paradigms that rely on pre-defined, isolated stimuli, spectral novelty is computed in relation to the surrounding acoustic context. This means that the algorithm dynamically evaluates whether a sound deviates from the local sound environment. A sound may be classified as novel in one context but not in another, depending on its spectral contrast with the preceding acoustic input. This context sensitivity allows for a more ecologically valid identification of attention-capturing events, as it mirrors the perceptual mechanisms by which human listeners extract meaningful signals from background noise.

By leveraging this context-aware detection of acoustic change, we move beyond discrete stimulus presentations toward a more naturalistic framework for studying auditory attention. The three-phase model of distraction (Escera et al., 2000; Getzmann et al., 2024; Wetzel & Schröger, 2014), typically investigated in tightly controlled laboratory settings, can thus be extended to complex real-world environments. Here, attentional shifts and reorienting responses may be shaped by factors such as habituation, cognitive load, and environmental complexity (Brockhoff et al., 2023; Gygi & Shafiro, 2011; Lavie, 2005; Woods & Elmasian, 1986). The successful application of spectral novelty in ERP research not only enhances ecological validity but also offers a promising tool for investigating dynamic attention in everyday auditory scenes.

### 4.6 Limitations and Future Directions

While our study provides valuable insights into auditory attention in real-world settings, certain methodological aspects warrant consideration. First, although spectral novelty served as a robust marker of auditory salience, it does not capture other factors such as semantic relevance or emotional valence, which can also strongly influence attention allocation (Asutay & Västfjäll, 2012; Debnath & Wetzel, 2022; Holtze, Jaeger, Debener, Adilŏglu, & Mirkovic, 2021; Kjellberg et al., 1996; Lavie, 2005; Roye, Jacobsen, & Schröger, 2013). Second, the fixed order of conditions may have introduced habituation effects, as participants were always exposed to the same sequence of listening modes. Counterbalancing condition order in future studies would help disentangle potential order effects from true condition-related differences.

Furthermore, while we observed clear P3a responses to acoustic novelty, the absence of strong condition effects suggests that our task manipulations may not have been sufficiently distinct to drive measurable differences in ERP amplitude. Refining the contrast between passive and active listening may help clarify how task demands shape auditory distraction. Finally, future studies should consider investigating individual variability in noise sensitivity, as subtle differences in attentional engagement may be masked in group-level analyses of P3a and RON components.

## 5 Conclusion

In conclusion, our study provides compelling evidence that spectral novelty serves as a reliable and ecologically valid trigger of attentional processing in naturalistic soundscapes. Across a large dataset and diverse listening contexts, we found that higher novelty consistently elicited strong P3a responses, demonstrating robust neural signatures of attentional capture even when participants were engaged in unrelated tasks. Importantly, this neural response was accompanied by measurable behavioral slowing, confirming that these sound events were not only registered by the brain but also disrupted ongoing performance.

These findings highlight the value of spectral novelty detection as a powerful tool for identifying cognitively relevant sound events in real-world environments, moving beyond traditional stimulus designs. The P3a emerged as a particularly stable marker, showing consistent morphology and amplitude across conditions, positioning it as a key component for studying auditory distraction outside the lab. While listening mode did not strongly influence ERP amplitudes, this likely reflects the adaptive nature of auditory attention rather than a lack of engagement. Similarly, the absence of a correlation with noise sensitivity underscores the idea that neural responses to distraction are more influenced by moment-to-moment context than by trait-level sensitivity.

Altogether, our results underscore that the brain remains highly responsive to acoustic novelty in real-world settings, both neurally and behaviorally and establish spectral novelty detection as a promising approach for future research on attention, cognition, and distraction in everyday life.

## 6 Data and Code Availability

The stimuli, analysis scripts and data to reproduce the findings of this paper can be found at: https://zenodo.org/records/15182196

## 7 Author Contributions

SK and MB conceptualized the experiment. SK performed data acquisition, conducted the analyses, and wrote the manuscript. MB and TH provided critical revisions. TH contributed the concept and implementation of the spectral novelty detection algorithm by adapting a public Python script for use in MATLAB, which formed the basis for all novelty-based analyses. All authors approved the final version of the manuscript and agreed to be accountable for the work.

## 8 Funding

This work was funded by the Deutsche Forschungsgemeinschaft (DFG, German Research Foundation) under the Emmy-Noether program–BL 1591/1-1–Project ID 411333557.

## 9 Acknowledgements

The authors would like to thank Daniel Küppers for his kind help with the typing speed analysis. Further, we would like to thank all members of the NELI group for their support and guidance. The authors would also like to thank Negar Dadkhah and Amrah Gasimli for their annotation of the soundscape and the Friedrich Ebert Foundation, which always provides helpful support to SK.

## 10 Conflict of Interest

The authors declare that the research was conducted in the absence of any commercial or financial relationships that could be construed as a potential conflict of interest.

## 11 Supplementary Material

**Figure 9:**
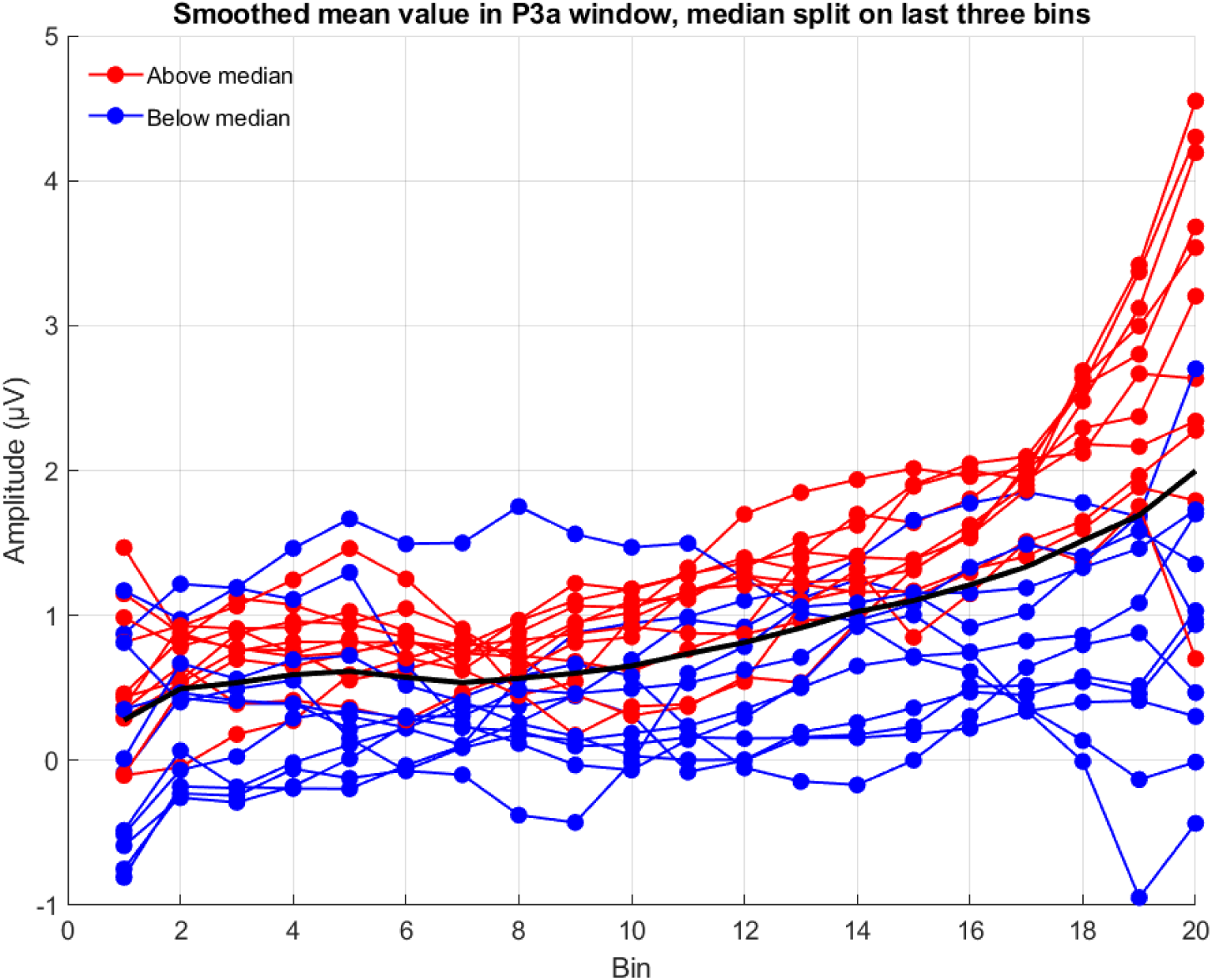
Smoothed mean EEG amplitude in the P3a window across spectral novelty bins and averaged over all conditions, split by median amplitude in the highest novelty bins. Red lines represent participants with amplitudes above the median, blue lines represent participants below the median, and the black line shows the grand average. This figure illustrates the overall trend of increasing EEG amplitude with spectral novelty, with inter-individual variability in the magnitude of this effect.

**Figure 10:**
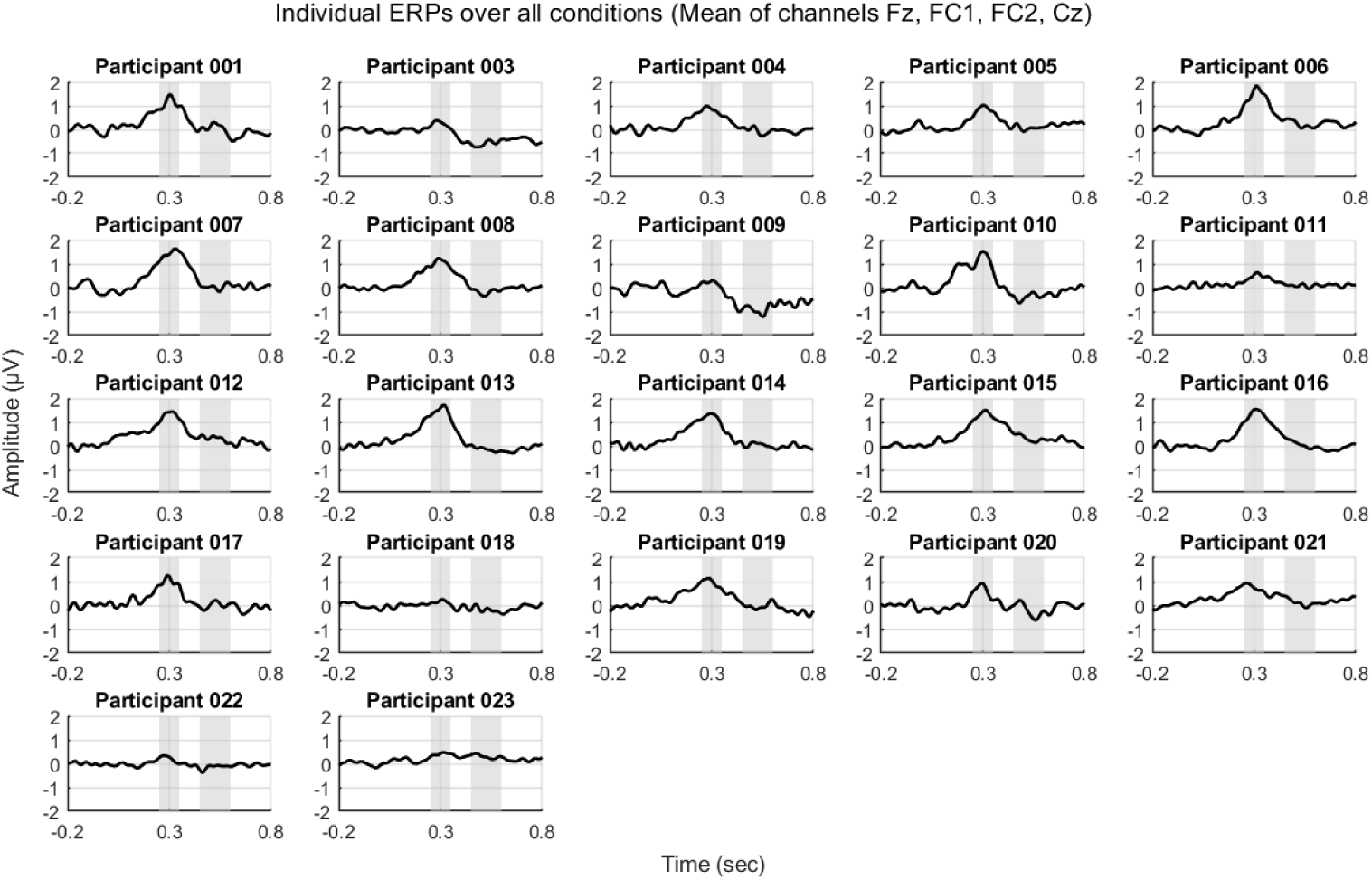
Individual ERPs for each participant, averaged across all conditions at selected frontocentral electrodes (Fz, FC1, FC2, and Cz). Shaded regions indicate the time windows of interest: P3a (250–350 ms, light gray) and Reorienting Negativity (RON, 450–600 ms, dark gray).

## Footnotes

1 https://www.youtube.com/watch?v=Le_g4s6KloU, (accessed 01.07.22)

2 for further information see: https://www.zooniverse.org/projects/courtaulddigital/world-architecture-unlocked/about/research

3 https://github.com/labstreaminglayer/liblsl-Matlab, v1.14.0.

4 https://github.com/labstreaminglayer/App-Input, v1.15.0.

5 https://github.com/labstreaminglayer/App-LabRecorder, v1.14.0.

## Notes

https://zenodo.org/records/15182196

### Competing Interest Statement

The authors have declared no competing interest.

